# Experimental and computational framework for a dynamic protein atlas of human cell division

**DOI:** 10.1101/227751

**Authors:** Yin Cai, M. Julius Hossain, Jean-Karim Hériché, Antonio Z. Politi, Nike Walther, Birgit Koch, Malte Wachsmuth, Bianca Nijmeijer, Moritz Kueblbeck, Marina Martinic Kavur, Rene Ladurner, Stephanie Alexander, Jan-Michael Peters, Jan Ellenberg

## Abstract

Essential biological functions, such as mitosis, require tight coordination of hundreds of proteins in space and time. Localization, timing of interactions and changes in cellular structure are all crucial to ensure correct assembly, function and regulation of protein complexes^1-4^. Live cell imaging can reveal protein distributions and dynamics but experimental and theoretical challenges prevented its use to produce quantitative data and a model of mitosis that comprehensively integrates information and enables analysis of the dynamic interactions between the molecular parts of the mitotic machinery within changing cellular boundaries.

To address this, we generated a 4D image data-driven, canonical model of the morphological changes during mitotic progression of human cells. We used this model to integrate dynamic 3D concentration data of many fluorescently knocked-in mitotic proteins, imaged by fluorescence correlation spectroscopy-calibrated microscopy^5^. The approach taken here in the context of the MitoSys consortium to generate a dynamic protein atlas of human cell division is generic. It can be applied to systematically map and mine dynamic protein localization networks that drive cell division in different cell types and can be conceptually transferred to other cellular functions.

## RESULTS

To generate standardized, quantitative data on the dynamic 3D localization of mitotic proteins, we imaged HeLa cell lines, in which such proteins were fluorescently labeled by editing the corresponding genomic locus ^6^. For each protein, the cell and chromosome volumes were recorded in separate channels as spatio-temporal landmarks. We recorded mitosis in high throughput by detecting the beginning of cell division (prophase) in low resolution images of the chromosomes, imaging the protein of interest by high resolution 3D confocal microscopy until division was completed (**Fig. 1a**) and then calibrating the signal by fluorescence correlation spectroscopy (FCS)^7^ in six nuclear/cytoplasmic positions (**Fig. 1a**). Calibration allowed us to convert 3D protein fluorescence movies to time-resolved protein concentration distribution maps (see Methods; **Fig. 1b,c)**. Using this automated experimental pipeline, we acquired a pilot dataset for 28 proteins, most of which were homozygously tagged with EGFP by zinc finger nucleases^8^ or CRISPR-Cas9 nickase^9^ mediated genome editing, while for some genes stable integration of cDNAs or bacterial artificial chromosomes^10^ had to be used (see Methods; **Supplemental Data Table 1**). The time-resolved 3D distribution was recorded for 18 dividing cells per protein on average (**Fig. 2a**), giving us a sufficiently large dataset to develop and test our computational framework.

**Figure 1.**
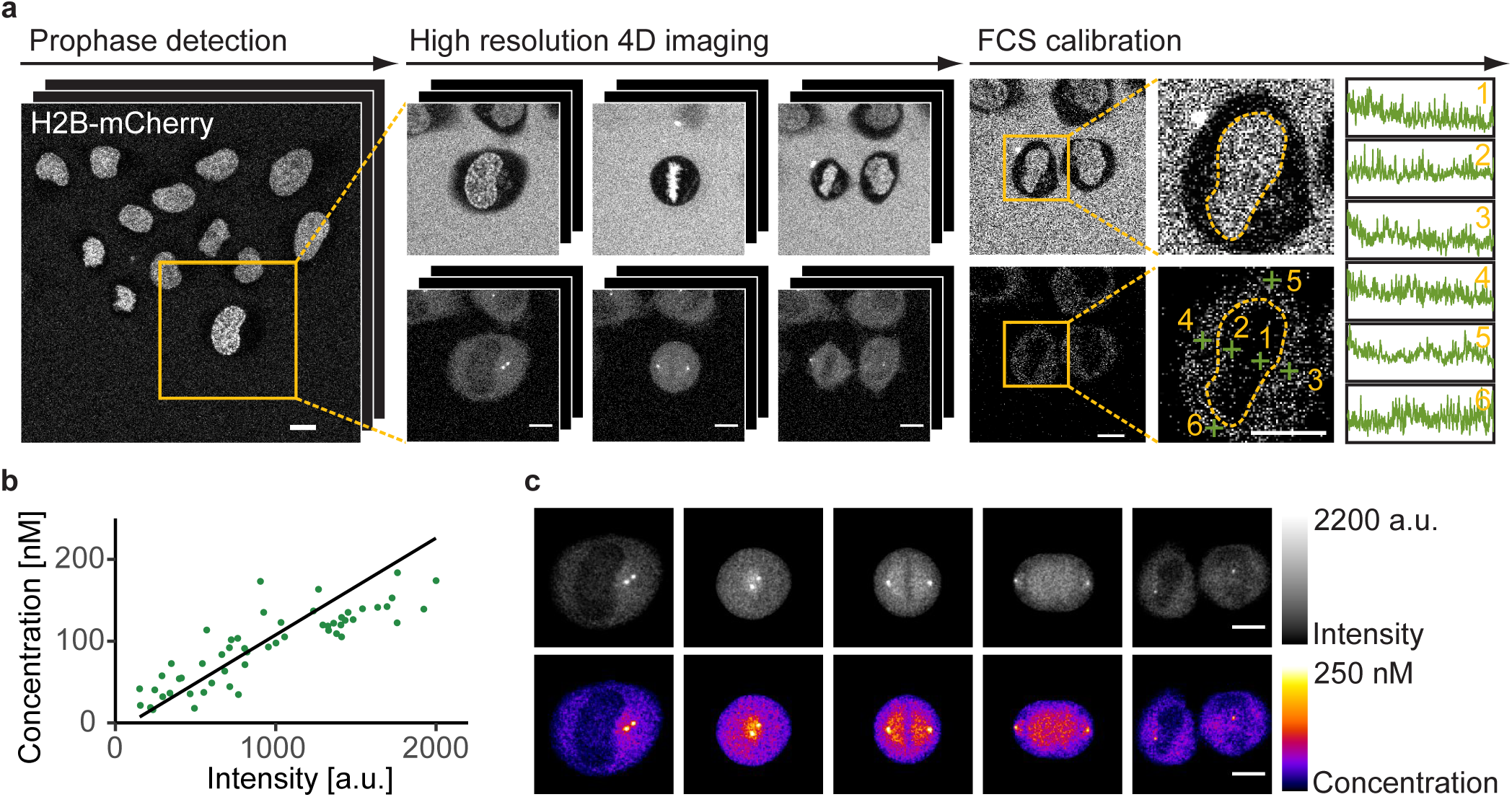
Quantitative imaging of mitotic proteins. (a) Automatic calibrated 3D live confocal imaging pipeline. Cells in prophase were identified by online classification, imaged through mitosis in the landmarks and protein of interest channels, and measured by FCS at selected positions. (b) The local protein concentrations determined by FCS fitting linearly correlate with the background subtracted image intensities at the corresponding positions (shown are data acquired on the same day). (c) Example cell showing concentration map resulting from FCS-based intensity calibration (mean z-projection). Scale bar: 10 μm. Data shown in (a)-(c) is for H2B-mCherry mNEDD1-LAP (EGFP) and is representative of *n* = 92 independent experiments performed with 28 different cell lines.

**Figure 2.**
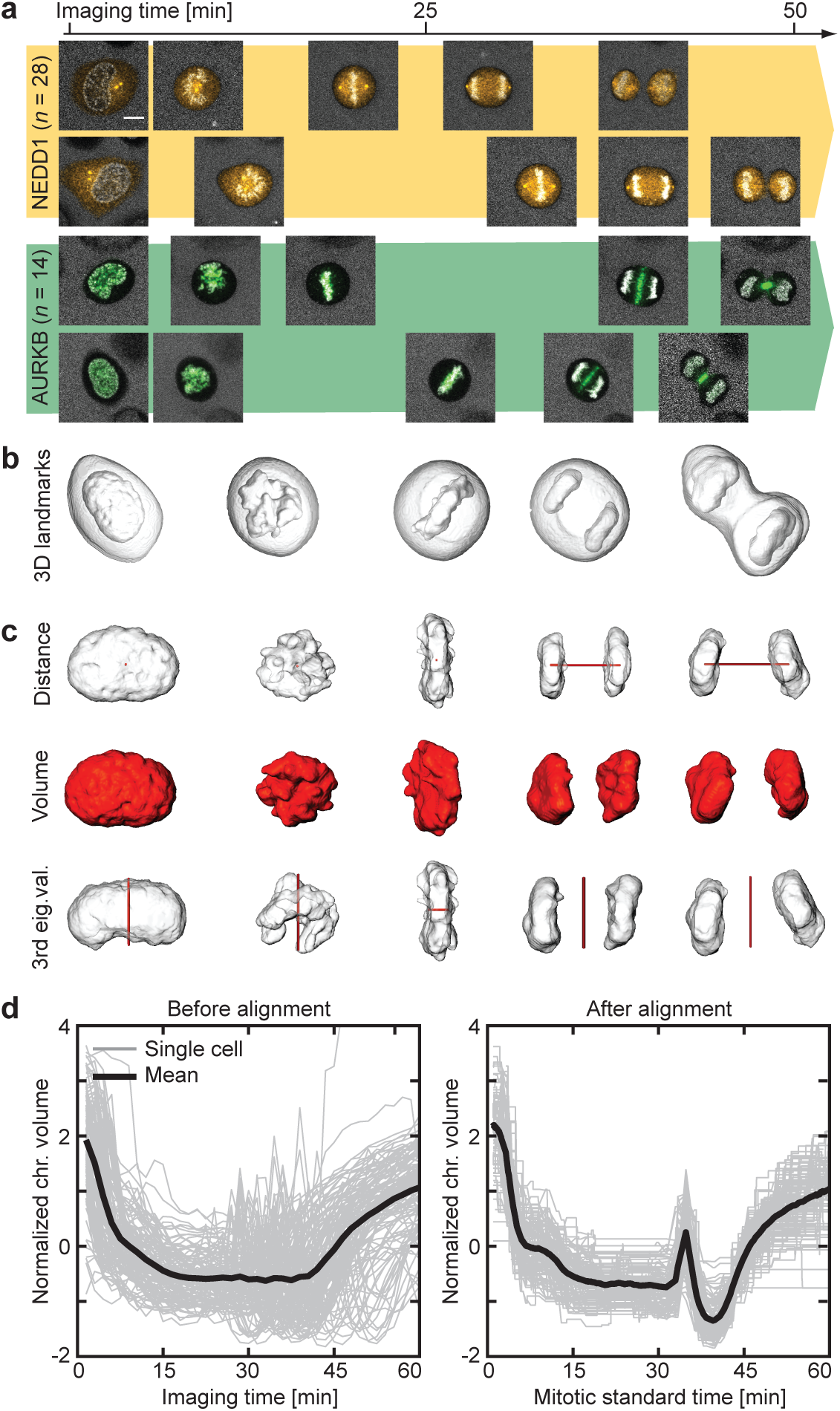
Modeling of mitotic standard time. (a) Individual cells have different mitotic spatio-temporal dynamics. Scale bar: 10 μm. (b) Cellular and chromosomal volumes were segmented from the landmarks channel. (c) Three morphological features (in red) were extracted from the chromosomal volume. (d) Mitotic standard time was generated in the feature space by multiple sequence alignment visualized here in the feature dimension describing chromosomal volume. Shown is the alignment of *n* = 132 cells from 20 independent experiments.

Although cell division is a continuous process, traditionally, mitosis is divided into five stages: pro-, prometa-, meta-, ana- and telophase^11^. Except for nuclear envelope breakdown and chromosome segregation that mark the onset of prometaphase and anaphase respectively, the other stages are not separated by sharp kinetic boundaries. To align the varying kinetic data from different cells (**Fig. 2a**), we first defined a “mitotic standard time” based on changes in chromosome structure. Chromosome boundaries of all imaged mitoses were automatically segmented in 3D using the landmark channel (see Methods; **Fig. 2b**, **Extended Data Fig. 1a,b)**. Three geometric features were extracted from the segmented data: the distance between the two segregated chromosome masses, the total chromosome volume and the third eigenvalue of the chromosome mass (**Fig. 2c**). Each mitosis movie could thus be represented as a six-dimensional vector sequence of these parameters and their first derivative indicating kinetic transitions. We used a modified Barton-Sternberg algorithm with multidimensional dynamic time warping to align the vector sequences and construct a mitotic standard time reference (see Methods; Fig 2d, **Extended Data Fig. 1c,d)**. To discretize major transitions in chromosome structure during mitosis, we detected local maxima in the second derivative of the average feature sequences (**Extended Data Fig. 2a-c**). This automatically distinguished 20 mitotic stages, which we used to annotate the experimentally sampled time points of individual HeLa cells throughout the study (**Extended Data Fig. 2d**). The same approach could align a different human cell type, U2OS, using the same landmarks, and conserved the nature of the mitotic transitions (**Extended Data Fig. 3**) validating the generality of the approach.

This alignment allowed to objectively map all cell images to a standard time reference for averaging. To enable visualization, interactive navigation and analysis of all imaged protein distributions, we next computed the canonical geometry from late prometaphase (stage 7) to cytokinesis (stage 20), where little deviation from rotational symmetry around the division axis occurs. The canonical geometry model was reconstructed from the average geometry of several hundred cells spatially registered for each mitotic standard stage (see Methods; **Extended Data Fig. 4**). Evolution of this mitotic standard geometry over the mitotic standard time defines the 4D canonical mitotic cell model, enabling us to register all recorded cell divisions in space and time based on their landmark channels. For each protein, we mapped each 3D stack to the corresponding mitotic standard stage (**Fig. 3a**) and computed 4D concentration maps representing the average behavior of each mitotic protein. Maps of many proteins can then be freely combined (**Fig. 3b**), to compare their localization patterns, dynamics and abundance and provide intuitive navigation of the integrated data set as illustrated in our web-based interactive mitotic cell atlas (www.mitocheck.org/mitotic_cell_atlas).

**Figure 3.**
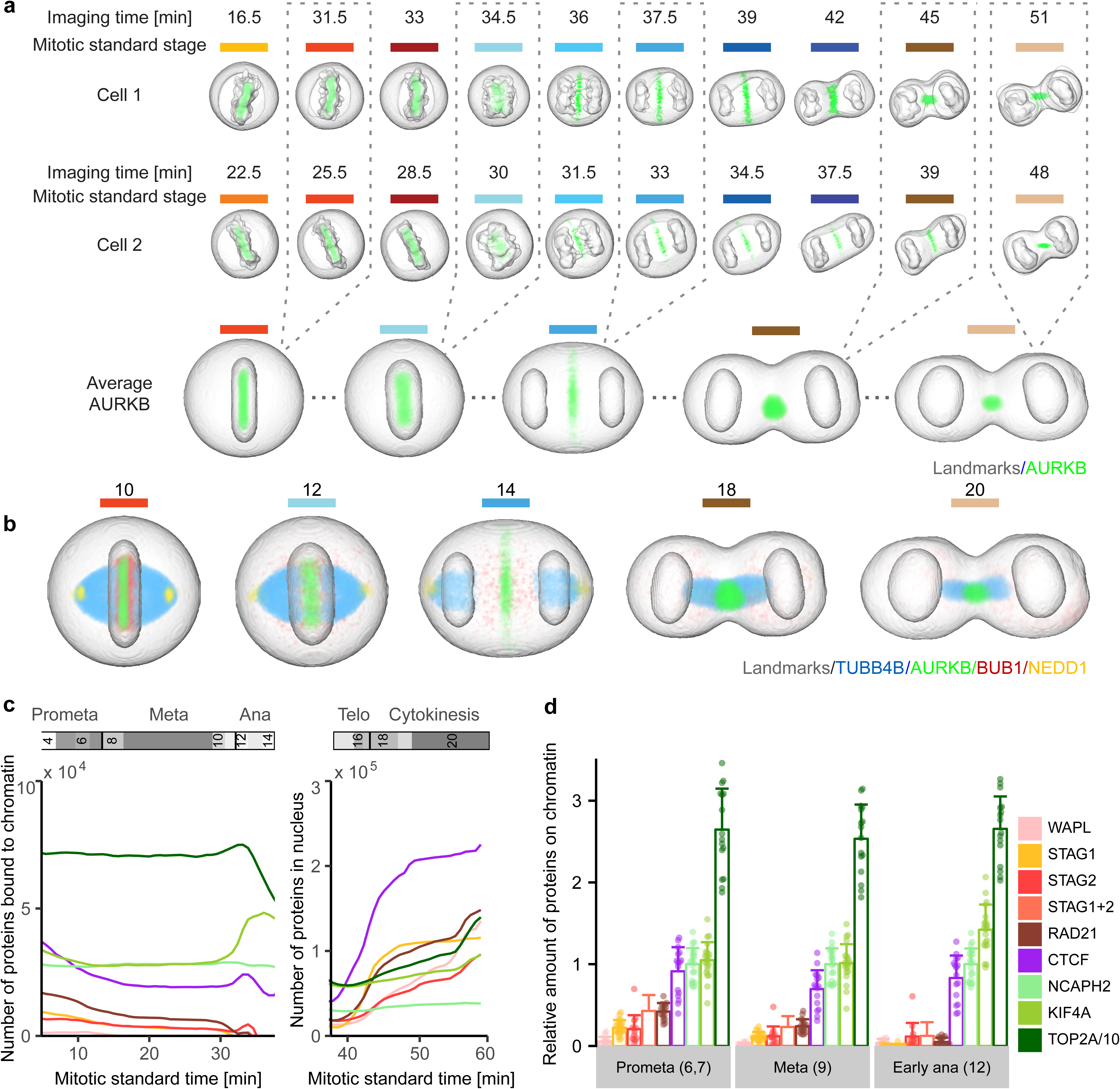
Visualization of 4D protein distribution maps. (a) Through averaging of a large number of cells, models were generated for all mitotic standard stages with symmetrical geometries. Example image sequences were registered to the standard space of the corresponding mitotic standard stage. A distribution map over time was then generated for each protein by averaging through multiple cells. Colored lines indicate mitotic stages. (b) Average distributions of four proteins are displayed in different mitotic stages. (c) Amount of chromatin-bound and nuclear molecules for eight chromatin remodelers. (d) Fraction of chromatin bound proteins relative to NCAPH2. Shown are the single cell values (dots) and the mean and standard deviation. The sum of STAG1 and STAG2 (STAG1+2) was calculated from the mean and standard deviation of STAG1 and STAG2 data. In (c) and (d), TOP2A has been scaled down by a factor 10 for visualization. Note: reported numbers represent monomers, dimers (e.g. TOP2A) would result in a 50% reduced abundance of complexes.

To illustrate the power of integrated data exploration for multiple proteins in the canonical model, we analyzed eight mitotic chromosome structure proteins (**Extended Data Fig. 5a,b)**. Plotting the total number of proteins found on mitotic chromosomes and in the daughter nuclei against the mitotic standard time allowed a quantitative comparison of protein dynamics (see Methods; **Fig. 3c,d)** which revealed that the amount of most chromosomal proteins bound to chromatin in metaphase is within the same order of magnitude (3,000 - 26,000 per nucleus), except for TOP2A which shows a 25 times higher abundance, potentially suggesting a structural rather than a purely enzymatic role. We found cohesins to slowly and progressively dissociate from chromatin in early mitosis (RAD21, STAG1, and STAG2), with a final more rapid release of approximately 3,000 remaining cohesin complexes before anaphase onset, indicating that no more than 100 cohesins are sufficient to connect the sister chromatids on an average human chromosome, mostly at the centromere (see Methods and ^12^). Interestingly, the cohesins bound to mitotic chromosomes consisted of equal amounts of two isoforms containing the HEAT repeat subunits STAG1 or STAG2 (**Fig. 3d**) contrasting with the situation in interphase nuclei where STAG2-containing complexes dominated^13^. Furthermore, we observed that a significant amount of STAG2 (*p* < 0.025, paired Wilcoxon signed rank-test), but not of the kleisin subunit RAD21, rebound chromosomes in anaphase, suggesting a potential non-cohesive function of STAG2 during mitotic exit (**Fig. 3c,d**, **Extended Data Fig. 5b**). In contrast to the complete dissociation of most cohesins, about 17,000 molecules of the chromatin organizer CTCF remained associated with the genome throughout mitosis^14^, consistent with a “bookkeeping” mechanism of interphase chromatin architecture. Once chromosome segregation was initiated, KIF4A, TOP2A and CTCF further accumulated on chromatin in anaphase, suggesting a role in maximal arm shortening in anaphase^15^. During nuclear reformation, the cytoplasmic pool of mitotic chromosome proteins showed an ordered entry as well as decreasing rates of import. CTCF was reimported first with the highest rate (391 proteins/sec), followed by simultaneous import of the cohesin subunits STAG1 and RAD21 (352 and 239 proteins/sec, respectively), while STAG2 and WAPL enter the nucleus later and at a lower rate (69 and 89 proteins/sec, respectively). This shows that mitotic decondensation proceeds in the presence of CTCF and subsequently STAG1-containing cohesin complexes, but before WAPL and STAG2 are present. In addition to the chromosomal proteins, we also explored assembly of the nuclear pore complex (NPC) during late anaphase (**Extended Data Fig. 5c**). Consistent with previous observations^16,17^, we found that cytoplasmic ring components (NUP107, NUP214) assembled early, but surprisingly found that nuclear basket as well as cytoplasmic filament proteins (TPR and RANBP2) assembled only much later, at a time when import of CTCF was already completed. This suggests that nuclear and cytoplasmic filaments of the NPC are not required for the rapid import of nuclear proteins needed for re-establishing the interphase genome architecture (**Extended Data Fig. 5d**).

To comprehensively investigate which proteins work together where and when inside the cell, we transformed their spatial distribution into numerical features. To this end, we used a segmentation-free approach based on a speeded-up robust features (SURF) detector^18^ to extract so-called interest point clusters (see Methods; **Fig. 4a**, **Extended Data Fig. 6a,b)** transforming each 3D movie into a sequence of 100-dimensional feature vectors.

**Figure 4.**
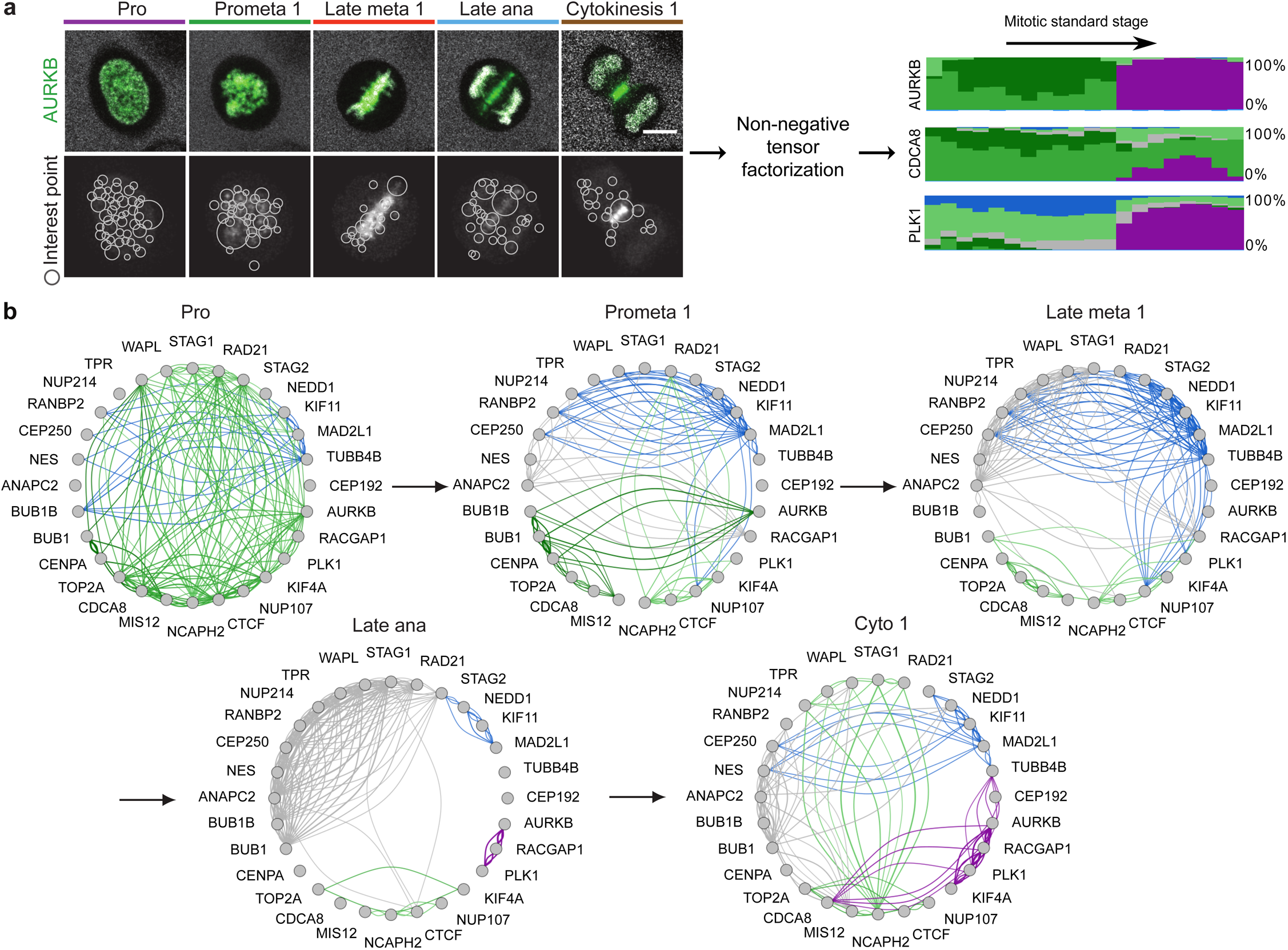
Identification of dynamic protein clusters. (a) SURF interest points were detected and assigned to one of 100 clusters of similar interest points. Non-negative factorization of the data tensor of 28 proteins × features × mitotic stages produced a non-negative tensor of reduced dimension whose entries can be interpreted as the fraction of protein belonging to each cluster over time (right panel, each cluster is represented by a different colour and the height of a coloured bar at a given mitotic stage represents the fraction of the protein in the corresponding cluster at this stage). Scale bar: 10 μm. (b) Dynamic multi-graph of protein co-localization, shown for 5 stages. Each edge colour corresponds to a localization cluster as in (a) and the edge thickness corresponds to the product of the linked genes fractions in the corresponding cluster and can be loosely interpreted as a probability of interactions.

By averaging the feature vectors of all images of the same protein and mitotic standard stage, the dynamic distribution of all proteins in our dataset could then be represented as a third order tensor of size 28 × 100 × 20 (proteins × features × stages). Soft clustering with non-negative tensor factorization (NTF) detected seven clusters of dynamic protein localization patterns (see Methods, **Fig. 4a**, **Extended Data Fig. 6c**). The identity of the proteins in each pattern revealed a striking correspondence between these statistically defined clusters and major mitotic organelles and structures (e.g. CENPA identifies centromeres/kinetochores, RACGAP1 identifies the midplane in late mitotic stages, **Extended Data Fig. 7**), validating our unsupervised approach. Since our clustering assigns the fraction of a mitotic protein to each pattern over time, it reliably deals with promiscuous proteins present in multiple sub-cellular structures. Linking proteins with similar patterns at each time point allowed us to derive a dynamic multigraph, which showed the dynamic protein co-localization network highlighting the activities of different compartments over time and allowing us to predict where and when proteins most likely interact (**Fig. 4b**). Results from the above clustering can be used to generate hypotheses that can be visualized in the canonical cell model. For instance, the temporal evolution of the mitotic kinase AURKB and its regulator CDCA8 (aka Borealin), predicts that the two proteins relocate to the midplane (**Fig. 4a**, purple in right panel) in different proportions and with different kinetics. Since the two proteins are known to be present in a 1:1 ratio in the chromosome passenger complex^19^, this observation suggested that a fraction of AURKB in the midplane is not part of the complex. Exploring the 4D localization of CDCA8 and AURKB in the mitotic cell atlas (**Extended Data Fig. 8a,b)** indeed revealed that while these two proteins partially colocalize at the midbody, AURKB shows an additional localization in an equatorial cortical ring that contracts as mitosis progresses. This novel localization of the mitotic kinase AURKB suggests that it is an integral part of the contractile cytokinetic ring. While unexpected, this observation is consistent with AURKB’s function in cytokinesis^20-22^. This raises the very interesting possibility that the midplane and cytokinetic ring pools of AURKB have different functions for central spindle architecture and cytokinesis respectively during mitotic exit.

Since our clustering of dynamic localization patterns does not a priori yield pure sub-cellular compartments as defined ultrastructurally or by fractionation, we developed a supervised machine learning approach to define subcellular structures based on known resident proteins of six compartments/organelles relevant for mitosis: chromosomes, nuclear envelope, kinetochores, spindle, centrosomes and midbody (see Methods). Using the interest point cluster features as input, we trained a multivariate linear regression model that could assign the amounts of a protein of interest present in each of the six reference compartments (**Extended Data Fig. 8c,d)**. This allowed us to quantitatively compare the subcellular fluxes between these compartments for all proteins (**Extended Data Fig. 9**).

Mining this data is powerful to dissect dynamic multimolecular events inside living cells such as the assembly or disassembly of organelles. As an example, we calculated the number of all imaged proteins localized to kinetochores, to investigate the disassembly of this large supramolecular complex essential for cell division. The data allowed us to determine that kinetochore disassembly starts in early metaphase with dissociation of BUB1B and PLK1 followed in late metaphase by BUB1, AURKB, MIS12 and CDCA8 (**Extended Data Fig. 8d**). In addition, this analysis showed that the stoichiometry of these proteins prior to disassembly differs up to six-fold and that their maximal dissociation rates span over an order of magnitude ranging from 17 to 173 molecules per second. The predicted number of ~420 CENPA molecules per kinetochore was consistent with data from biochemical methods^23,24^ and the predicted disassembly order was consistent with reports of late dissociation of MIS12^25^ (**Extended Data Fig. 8e**).

Our automatic assignment of protein amounts to cellular organelles, thus allowed us for the first time to determine the exact timing, stoichiometry and dissociation rates for multiple mitotic proteins that reside dynamically on kinetochores.

## DISCUSSION

With this study we provide an integrated experimental and computational framework to build a comprehensive and quantitative 4D model of the mitotic protein localization network in a dividing human cell. Our model provides a standardized yet dynamic spatio-temporal reference system for the mitotic cell that can be used to integrate quantitative information on any number of protein distributions sampled in thousands of different single cell experiments. Using a pilot data set, we illustrate the power of this model by mining the data to automatically define dynamic localization patterns to subcellular structures as well as predicting the order, stoichiometry and rates of assembly and disassembly of sub-cellular organelles. This quantitative information on protein localization in living cells provides greater insights into protein dynamics and interactions at relevant temporal resolutions and supports building simulations of mitotic processes. Our computational model underpins an interactive 4D atlas of the human mitotic cell, which allows the visualization of multiple protein dynamics with a spatial and temporal resolution and continuity that are currently very difficult or impossible to reach with multi-color live-cell imaging over the duration of mitosis. We illustrate with mitotic chromosome formation, kinetochore disassembly, NPC assembly and cytokinesis how the knowledge gained through the exploration and mining of the atlas data can be used to formulate new mechanistic hypotheses about the function of proteins inside the cell. The concept of standardizing the spatio-temporal cellular context for analyzing dynamic protein distributions in order to understand cellular processes as presented here is generic and we envision its adaptation to other essential biological functions such as cell migration or cell differentiation.

## ACKNOWLEDGEMENTS

We thank T. Hyman and I. Poser for donating multiple mouse BAC protein-GFP cell lines and T. Hirota for giving us the EGFP-CENPA cell line. The automatic imaging would not have been possible without the generosity of R. Höfler and D. W. Gerlich who developed the Micronaut software. We thank all members of the Ellenberg and Peters laboratories for support, especially M. Isokane, M. J. Roberti, J. Mergenthaler, S. Otsuka, W. Tang and D. Cisneros for generating cell lines, reagents and constructs. We thank A. Callegari for supporting the U2OS data generation. We also thank W. Huber, B. Fischer, B. Klaus and L. P. Coelho for discussions, the EMBL mechanical and electronic workshop, the EMBL advanced light microscopy facility and the EMBL flow cytometry core facility for their excellent support. This study has hugely benefited from the collaboration with Carl ZEISS Jena, especially with T. Ohrt. The work was supported by grants from EU-FP7-MitoSys (Grant Agreement 241548) to J.E. and J.M.P., the EU-FP7-SystemsMicroscopy NoE (Grant Agreement 258068) and EU-H2020-iNEXT (Grant Agreement 653706) both to J.E., as well as by the European Molecular Biology Laboratory (Y.C., M.J.H., J.K.H., A.Z.P., N.W., B.K., M.W., B.N., M.K., S.A., J.E.). Y.C. and N.W. were also supported by the EMBL International PhD Programme (EIPP). Research in the laboratory of J.M.P. was further supported by Boehringer Ingelheim, the Austrian Science Fund (FWF special research program SFB F34 “Chromosome Dynamics” and Wittgenstein award Z196-B20), the Austrian Research Promotion Agency (Headquarter grants FFG-834223 and FFG-852936) and the European Research Council (ERC) under the European Union’s Horizon 2020 research and innovation programme (Grant Agreement 693949).

## AUTHOR CONTRIBUTIONS

B.K, B.N., M.K., R.L., M.M. and N.W constructed and validated GFP knock-in cell lines. A.Z.P. with the help of Y.C and M.W. developed the imaging pipeline. Y.C., A.Z.P., and N.W. performed the calibrated live cell imaging. Y.C., M.J.H., J.K.H., and A.Z.P. developed the computational analysis pipeline. Y.C., J.K.H., M.J.H., A.Z.P., S.A. and J.E. wrote the manuscript. J.M.P. coordinated the MitoSys consortium and supervised part of the work. J.E. supervised the work overall and originally conceived the project.

## METHODS

### Cell culture

HeLa Kyoto cells (RRID: CVCL_1922) were a kind gift from Pr Narumiya, Kyoto University. These cells were authenticated by whole genome sequencing. HeLa Kyoto cells were cultured in high glucose Dulbecco's modified Eagle's medium (DMEM; Life Technologies) supplemented with 10% (v/v) fetal bovine serum (FBS), 100 units/ml penicillin, 0.1 mg/ml streptomycin, 2 mM Glutamine and 1 mM (v/v) Sodium pyruvate at 37 °C and 5% CO_2_. Depending on the genetic modification, one or more of the following antibiotics were supplied to the culture at the stated final concentration: Geneticin (Life Technologies) 500 μg/ml, Hygromycin B (Invitrogen) 200 μg/ml or Puromycin (Invitrogen or Calbiochem) 0.5 μg/ml. Once the cells reached 80-90% confluence, they were passaged and only a fraction of the cells were cultured in a fresh dish. U2OS cells were obtained from the ATCC (HTB-96) and were not further authenticated. The U2OS cells were cultured in McCoy's 5A medium (Sigma-Aldrich) supplemented with 10% (v/v) FBS, 100 units/ml penicillin, 0.1 mg/ml streptomycin, 2 mM Glutamine, 1 mM (v/v) Sodium pyruvate, and 1% (v/v) MEM non-essential amino acids (Gibco). All cells tested negative for mycoplasma contamination.

### Cell modification

HeLa Kyoto cells were used for genetic modifications and imaging. HeLa Kyoto cells are hypotriploid with on average 64 chromosomes, thus during mitosis the cells have on average 64*2 = 128 kinetochores^12^. The cell lines have been generated for this project or previously^16,26-32^ are listed in **Supplementary Table 1** with their providers indicated. Several cell lines were generated within this project as follows: The cell lines expressing PLK1-mEGFP, CEP192-mEGFP and mEGFP-NUP107 were generated using the Zinc finger nuclease (ZFN) pipeline as in^29^. Zinc finger nucleases were purchased from Sigma-Aldrich with DNA-binding sequences listed in **Supplementary Table 2**. The other genome-edited cell lines were generated using the CRISPR/Cas9 system^9^ based on the paired Cas9 nickase approach. For these cell lines, both gRNAs (**Supplementary Table 2**) and the donor plasmid were designed based on ENSEMBL release 75 and transfected together into HeLa Kyoto cells with jetPrime (Polyplus) according to the manufacturer’s instruction. A single clone was selected using our previously developed validation pipeline^6,29^. For 4 out of 20 genome-edited cell lines (BUB1B-EGFP, TPR-mEGFP, mEGFP-NUP107 and CEP192-mEGFP) we detected in the Western blots (anti-GFP, Roche cat#11814460001) a band of the size of free GFP. Therefore, for these cell lines, the total free amount may be overestimated. In order to label the chromosomal volume, an H2B-mCherry^33^ cDNA was transfected into some genome-edited cell lines with Fugene6 (Promega) according to the manufacturer’s instructions. The pmEGFP2-N1-NES construct was generated by sub-cloning two tandem repeats of mEGFP (mEGFP2)^34^ and the NES of MAPKK (NLVDLQKKLEELELDEQQ) into the pEGFP-N1 vector (Clontech Laboratories). The pmEGFP2-N1-NES construct was transfected into HeLa Kyoto cells and cells with stable expression were selected by culturing with the appropriate antibiotics. Single cells or a cell population with the desired expression level were harvested for imaging by fluorescence activated cell sorting (FACS, performed by the EMBL Heidelberg facility).

### Calibrated fluorescence confocal microscopy

Confocal microscopy was performed on Zeiss LSM780, Confocor 3, laser scanning microscopes using 40x, NA 1.2 water DIC Plan-Apochromat objectives and the GaAsP detectors equipped with an incubation chamber (EMBL workshop). Cells were imaged at 37 °C in a CO_2_-independent medium (Life Technologies) fluorescently colored with 500 kDa Dextran (Life Technologies) -DY481XL (Dyomics) produced in house. Time-lapse imaging was performed using the ZEN 2012 software as well as in-house software applications (see ^5^ for a software description). The acquisition was supported by an in-house developed objective cap and a water pump, such that water drops were regularly supplied to the objective-sample interface. Before starting imaging, a number of positions were selected manually. During live cell imaging, the microscope determined the focus automatically by performing line-scan imaging of the reflection signal of the 633 nm laser. The vertical position of the glass bottom was determined as the position with the maximum reflection intensity, and used as reference for acquiring a volume of the specimen at a particular depth.

The imaging workflow was set-up using the VBA Zeiss Macro MyPiC (https://git.embl.de/grp-ellenberg/mypic). HeLa Kyoto cell lines with H2B-mCherry were imaged live using an excitation laser at 561 nm every 5 min for about 16 hours on the Zeiss 780 microscopy system. Three confocal planes were acquired at a resolution of 0.32 μm in *x* and *y* and 2.5 μm in *z*. Images were projected in *z* by taking the maximum intensity value. Images of the H2B signal were analyzed on the fly by the Micronaut software (Gerlich lab, IMBA, Vienna) using a support vector machine classifier that was trained beforehand (with the software CellCognition, http://www.cellcognition.org/) to distinguish between cells in interphase, prophase, mitosis (prometaphase till telophase) and artefacts (apoptosis, on the border of the imaging field, out of focus, too low expression). The classification score for the prophase, interpreted as the probability of a cell being in the class of interest, was output, and a pre-defined threshold was used to make a decision on whether imaging setups for mitotic cell acquisition should be activated. According to how different the sample's H2B-mCherry expression levels were from the training set, the threshold on class probability was set between 0.85 and 0.96. Once a prophase cell was found, it was then imaged using a different imaging setup. For our purpose, mitotic cells were imaged live every 90 seconds for 31 *z*-planes with a spatial resolution of 0.25 μm in *x-y* and 0.75 μm in *z* with a 488 nm laser (high expression of H2B-mCherry allowed it to be excited at 488 nm and produce adequate signal). For cells not expressing H2B-mCherry we used SiR-DNA to stain the chromatin (Spirochrome, final concentration 50 nM added 2 hours before imaging). Cells were imaged with the 633nm laser (3 confocal planes every 7.5 min at the same resolution as for H2B-mCherry) and processed as for H2B-mCherry. For U2OS cells, the chromatin was stained with 200 nM SiR-DNA. To increase the incorporation of SiR-DNA the imaging media of U2OS contained 1 μM Verapamil (Spirochrome).

The signal from the GaAsP detector was separated into three channels: GFP, varied from 490 to 552 nm depending on the expression level to avoid detection saturation; mCherry, 587 – 621 nm (**Extended Data Fig. 1a, top row**), and Dy-481XL, 622 – 695 nm (**Extended Data Fig. 1a, second row**). For SiR-DNA we used 622 – 695nm. Once the mitosis was recorded for 40 frames, a single-plane image was then acquired at 2.5 μm above the cover glass surface. Using an adaptive feedback microscopy Fiji macro (https://git.embl.de/grp-ellenberg/adaptive_feedback_mic_fiji), the image was thresholded using the method developed by^35^ and the object closest to the image center with a proper size was selected as one of the two daughter nuclei in the cell of interest. The segmented nuclear boundary was fitted with an ellipse. FCS measurements were performed with the 488 nm laser and the APD detector at 505 – 540 nm at two positions within, and four around, the nucleus with a distance of 2 μm to the ellipse boundary for 30 s each. A manual quality control was then performed. Videos of cells with no expression of the protein of interest, with wrongly selected FCS positions (e.g. outside of the cell) or without anaphase onset were excluded from further processing. A total of 499 cells were retained with an average of 18 cells per protein (ranging from 10 to 35 cells per protein).

### Segmentation of the landmarks

A fully automated computational pipeline was implemented in MATLAB (MATLAB R2017a, The MathWorks, Inc., Natick, Massachusetts, United States) to segment and track cells of interest and reconstruct chromosomal and cellular surfaces (https://git.embl.de/grp-ellenberg/mitotic_cell_atlas). The pipeline was composed of three major steps: segmentation of chromosomal volume, segmentation of cell volume and extraction of parameters out of the landmarks geometry. Chromosomal regions were segmented from the mCherry channel which had high H2B–mCherry signal and very low Dextran-Dy481XL intensity (**Extended Data Fig. 1a, top row**) or from the SiR-DNA channel which had no crosstalk from other channels. In order to perform isotropic 3D image processing, adjacent *x-y* planes were linearly interpolated along the *z* direction. A 3D Gaussian filter was applied to reduce the effects of noise. To detect chromosomal regions, the filtered image stack was binarized first using a multi-level thresholding method as described in^36^. In this approach, a global Otsu threshold^37^ was determined for the entire stack and the threshold was then adapted for each 2D slice, validated by the connectivity of binary components in 3D. Tiny connected components were removed from the binary image leaving only chromosomal components from all cells in the imaging field. All components were used as seeds for the detection of the cell boundary in a later stage. The connected chromosomal volume in the *x-y* center of the first frame identified the cell of interest due to the centering step in the imaging pipeline. The cell of interest was tracked sequentially through the entire image sequence using a nearest neighbor approach. At each time point, an event of chromosome segregation was also probed by analyzing the chromosomal volume around the tracked location. Once segregation was detected, both daughter nuclei were tracked in the subsequent frames (**Extended Data Fig. 1a, third row**).

The cell region was segmented from the Dy481XL channel showing the cell-free regions in high intensities and histone signal with low intensities (**Extended Data Fig. 1a, second row**). Upon interpolation and filtering as for the chromosomal segmentation, a ratio image was created by dividing the filtered image stack of the mCherry channel by that of the Dy481XL channel in order to diminish bleed-through signal from the H2B-mCherry channel. The ratio image stack was then binarized as described above. When using SiR-DNA there was no bleed-through in the Dy481XL channel when excited at 488 nm. In this case the Dy481XL was directly binarized. To separate individual cell regions, the previously detected nuclear seeds were used, considering the fact that each cell region can have only one or two chromosomal volumes. This was implemented by applying a marker-controlled watershed algorithm^38^. To obtain a better separation between touching cells, the algorithm was applied on the distance transformed image that made use of the geometric properties of the cell surface. The cell region of interest was defined by taking the connected region(s) containing the detected chromosomal volume(s) of interest (**Extended Data Fig. 1a, bottom row**).

The chromosomal mass at each time point was represented by its three orthogonal eigenvectors and associated eigenvalues where the eigenvector with the largest eigenvalue represented the longest elongated axis of the chromosomal volume. Metaphase frames were automatically detected based on the low value of the smallest eigenvalue of the chromosomal volume. Division axis for metaphase cells was then predicted by taking the eigenvector having the minimum eigenvalue. By definition, this vector is always orthogonal to the metaphase plate. Using the predicted axis in the first and last metaphase frame, axes for the remaining frames were propagated backwards and forwards for stages before and after metaphase, respectively, where the eigenvector with the smallest discrepancy in angle to the axis predicted for the adjacent frame was used (**Extended Data Fig. 1b**). For further analysis, the plane orthogonal to the division axis going through the centroid of both daughter nuclei was predicted as the midplane. Segmented landmarks were 3D reconstructed and visualized for quality control. Cells having few time points over or under segmented were reprocessed with different parameters or using the results of correctly segmented adjacent time points as constraint.

### Image processing and calibration

Image processing and calibration were performed according to^5^. Before each calibrated live cell confocal microscopy experiment, the focal volume was calibrated using a 10-50 nM solution of Alexa488 (Life Technologies), and single mEGFP brightness was calibrated by performing FCS measurements on HeLa Kyoto cells expressing mEGFP^29^. All FCS measurements were processed using Fluctuation Analyzer^39^. Autocorrelation functions of dye solutions were fitted using a one-component diffusion model with triplet-like blinking, and measurements of fluorophore-fused proteins were fitted using a two-component anomalous diffusion model with fluorescent protein-like blinking. The effective confocal volume was calculated from

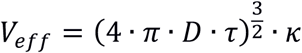

where *D* was the diffusion coefficient of the Alexa488, which is 464 μm^2^/s at 37 °C, and *κ* the structural parameter. The averaged time passing through the confocal volume *τ* and the structural parameter (typically between 4-7) were fitted to the ACF of Alexa488. The number of fluorescent molecules within a confocal volume was calculated by multiplying the fitted number of molecules (*N*) with correction factors for background and photobleaching^39^. As proteins might exist in complexes with multiple molecules, a count per molecule (CPM) value was used to correct the number of molecules. As reference, the CPM value of mEGFP was used as measured in the HeLa Kyoto cells expressing mEGFP where the mean value of all mEGFP measurements was taken. If the CPM of a measurement of a fusion protein of interest within a cell was larger than that of the mEGFP, the fitted number of molecules was corrected by multiplication with the ratio between the two. Finally, the local concentration of the measured protein was determined as the corrected number of molecules divided by the effective confocal volume. As quality control of the FCS measurements, we pre-defined thresholds and deleted data points with too low coefficient of variation *R*^2^ or APD counts or too high fitting *χ^2^* or bleaching or outlier CPM values.

The calibration of the image acquired with the GaAsP detector was based on the assumption of linear correlation between the local protein concentration and the EGFP intensity which we could verify (**Fig. 1b**). The averaged intensity of the GFP channel in cell-free areas was considered as background. For all measurement points the coefficient *ρ* between local protein concentration and background-corrected imaging intensity, mean filtered with a 9 × 9 pixels window to avoid noise, was calculated by performing a linear regression. The 3D protein concentration map was generated by multiplying the pixel intensities with the linear coefficient *ρ*. The protein number in each voxel was obtained by multiplying the concentration with the voxel volume. The absolute protein abundance could be calculated by summing up the map over the cell volume. After NEBD and before the nuclear envelope reforms, we estimated the number of proteins bound to chromatin by subtracting the cytoplasmic average concentration (representing the background concentration of proteins that freely diffuse between the cytoplasmic and chromosome volume) from the average protein concentration on the chromosome mask. Finally, to obtain the number of proteins, the concentration difference was multiplied by the number of voxels of the chromosome mask and the voxel volume.

To assess the accuracy of our quantitative measurements, we compared our data for nucleoporins (NUP107, NUP214, TPR and RANBP2) to expected numbers calculated from the known number of nuclear pores complexes (NPC) per cell and known protein stoichoimetry in each NPC. The HeLa Kyoto cell line used in this study has about 10,000 NPCs in interphase before nuclear envelope breakdown (NEBD)^16^. Assuming a nucleoporin (NUP) stoichiometry as reported ^40^ (32 NUPs/NPC for NUP107, TPR and RANBP2; 16 NUPs/NPC for NUP214), we can compute the number of NUPs present on the nuclear envelope (NE). Considering a free pool of nucleoporins in the cytoplasm that is included in our measurements, the ratio of our measurements over expected numbers on the NE should be greater than 1. We find that this ratio is on average 1.2 for all four NUPs, underlining the consistency of our measurements with established protein numbers by orthogonal methods.

### Modeling of the mitotic standard time

The mitotic standard time was modeled in a six-dimensional feature space using three morphological features of the chromosomal volume: the distance between the two daughter nuclei, the total volume and the third eigenvalue (**Fig. 2c**) and their first derivatives. The model was generated by aligning 132 mitotic image sequences using the Barton-Sternberg multiple sequence alignment algorithm (**Extended Data Fig. 1d**)^41^. The two sequences with the smallest distance to the average of all sequences were selected to initiate the alignment and each of the remaining sequences was then aligned to the average among all aligned sequences. The alignment was implemented as a modified multidimensional dynamic time warping^42^ where the total Euclidean distance over time between the pair of sequences was used as the objective of the optimization. The timeline of the averaged sequences was calculated as the mean of the alignment matrix as shown in **Extended Data Fig. 1c**. The Barton-Sternberg algorithm was terminated after four rounds as the standard deviation over time remained stable after 2 rounds (**Extended Data Fig. 1e**). The mitotic standard time was defined at a temporal resolution of 15 seconds by subsampling the averaged timeline. In order to find transitions in the mitotic standard time, the second derivative of the model at each time point for each feature dimension was calculated from

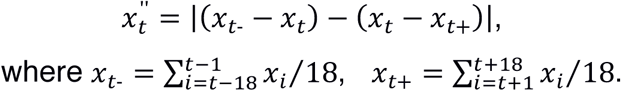

Peaks above a pre-defined threshold were selected across all dimensions as transitions (**Extended Data Fig. 2a**). In the later part of the model where the values of the second derivatives were generally low, small peaks were selected as additional transitions such that no stage between two transitions lasted longer than 12 minutes. Furthermore, transitions with lower values were deleted to ensure a minimum duration of 1.5 minutes for each stage (**Extended Data Fig. 2b,c)**.

This approach provides an objective way to discretize the mitotic standard time, which depends on the sampling and the number of cells used. Varying the number of cells sampled from our data identified between 19 and 21 stages with a median set of 20 mitotic stages, which we therefore used throughout the study. To check that these mitotic standard stages were biologically relevant we automatically selected the 3D image stack closest to the average feature values of each stage. Although the images picked in this way are from different cells, their automatically assigned sequential order reconstitutes a virtual mitosis with an error-free chronology (**Extended Data Fig. 2d**), in which all classically known mitotic transitions such as nuclear envelope breakdown (between stage 2 and 3), and anaphase onset (between stage 11 and 12), were correctly identified. Moreover, the method could identify previously hard-to-define stages such as the first formation of the metaphase plate in late prometaphase (between stage 7 and 8), and could differentiate between the different anaphase and telophase stages (stage 12 to 17). In addition, the kinetics of chromosome condensation is consistent with previous reports in different cell types ^15,36^ suggesting that the method could be applied to standardize the mitotic time in other cell types. To test this, we acquired a 4D image data set consisting of 43 U2OS cell divisions using the same imaging and landmarks approach. The same computational pipeline could indeed generate a mitotic standard time and mitotic standard stages for this cell line (**Extended Data Fig. 3**).

### Modeling of the canonical cell

To support spatial averaging, all cells assigned to the same standard mitotic stage were registered into a common reference coordinate system to give them the same location and orientation. To this end, a virtual coordinate system was defined with its origin at the center of a volume chosen large enough to accommodate all cells after registration. Landmarks (i.e. the cell boundary and chromosomal volumes) were then registered to the virtual coordinate system by applying a transformation function involving translation and rotation in 3D. This transformation function was estimated such that the predicted cell axis was aligned with the *x* axis in the virtual coordinate system. This transformation was applied to both landmarks to preserve their interrelationship in the registered image stacks as shown in **Extended Data Fig. 4a,b**. Bicubic interpolation was used when applying the transformation^43^.

Registered landmarks were subsequently represented using a cylindrical coordinate system that transforms 3D coordinates into radial distances providing greater flexibility in shape analysis. To this end, we converted landmarks in each plane along the *z* axis and along the predicted cell axis to polar coordinates in which object boundaries are represented by their radial distances from the object centroid (**Extended Data Fig. 4c**). As the centroids were aligned on the *z* axis, the cylindrical representation was formed by concatenating into a vector the polar representations for all planes (**Extended Data Fig. 4d**). After chromosome segregation, two separate cylindrical representations were used to encode each of the two daughter nuclei. In this case, the cylindrical axis of each chromosome passes through the centroid of that chromosomal volume.

The standard mitotic space represented by the averaged landmarks was computed in three steps. In the first step, the cylindrical coordinate vectors were averaged separately for each landmark across all cells within each standard mitotic stage (**Extended Data Fig. 4e**). The average vectors were then transformed back to a Cartesian coordinate system from which binary image stacks were generated. In a second step, to reconstruct the landmarks, the average volume of each landmark was obtained by combining two binary image stacks: one obtained using the *z* axis as the cylinder axis and the other using the cell axis as cylinder axis (**Extended Data Fig. 4f**). This combination involved first taking the intersection between the two binary images and then extending it until the average volume of all the cells belonging to the mitotic stage being processed was reached (**Extended Data Fig. 4f**). Because multiple frames of a cell could be assigned to the same mitotic standard stage, cells could have unequal contributions to each stage with some cells represented more than others at a given stage. To ensure uniform contribution from each cell towards the average mitotic space, in the third step, for each given mitotic stage and for each cell, the frame that was most similar to the average shape obtained in step 2 was selected. These selected cells were then used to re-compute the average shape of the corresponding mitotic stage as described above. This final average shape was also used to calculate the standard deviation of all cells in the same mitotic stage. Average mitotic space and standard deviation were generated for all the stages (7-20) for each of the landmarks (examples in **Extended Data Fig. 4g,h)**.

### Generation of the protein density map

Standard mitotic spaces were used as reference to register and integrate protein distributions from many different cells to generate protein density maps (**Fig. 3a**). All calibrated protein concentration maps having the same protein in a given mitotic stage were registered first to the corresponding standard mitotic space using the predicted cell division axis. This transformed all individual protein image stacks to the same coordinate system. Bicubic interpolation^43^ was used during the rotation. Registered image stacks were then accumulated in the standard mitotic space. Pixels outside the segmented cell region and mapped outside the standard mitotic space were discarded. A protein density map was then created by averaging the accumulated intensities in the standard mitotic space (**Fig. 3a**). Density maps of all proteins for mitotic stages 7-20 were estimated in the same way and can be explored on http://www.mitocheck.org/mitotic_cell_atlas.

### Feature extraction of images

The protein *z*-stack concentration map was processed using a Gaussian filter (Matlab smooth3 function with a kernel of size [3 3 1] and standard deviation 0.65) followed by a maximum projection along the *z*-axis and normalization to the theoretical saturation intensity. SURF interest points^18^ were then detected on the image resized to a 0.063 μm resolution using three octaves each including four Haar wavelet filters at different sizes from 9-by-9 till 99-by-99 pixels ranging from about half a micrometer to more than six micrometers. Interest points were further selected such that most of the protein signals were counted in one of the interest points. Each of these interest points was then described by a numerical vector quantifying features in the following four categories: locations relative to the landmarks (four features), correlation to the H2B signal or the predicted midplane/midbody volume according to the localization of the interest point (one feature), flattened soft spin image features^44^ describing the intensity distribution within an interest point (30 features), and summarized uniform Local Binary Patterns (uLBP)^45^ describing the orientation of the signal (4 features).

5% of the cells were randomly selected and their interest points were used to construct a training set for identifying clusters of interest points with similar features (**Extended Data Fig. 6a**). All training interest points were first separated into 16 clusters by their localization feature and the ¼-level of their metric values. Interest points in clusters with a sufficient size were then further clustered based on the correlation features separated by pre-defined thresholds followed by a dbscan^46^ clustering for each sub-cluster in the reduced feature space covering 85% of the variance according to a principal component analysis^47^ on the uLBP and spin image features^44^. The final clustering step was performed only for the clusters with the highest contrast value in their location category based on the spin image features where the interest points were separated into homogeneous bright, structured bright and dim clusters by pre-defined threshold. The total number of clusters was not deterministic since the training set was randomly generated but eight rounds of clustering yielded between 87 and 100 clusters and a run with 100 clusters was used for further analysis. Interest points in the same cluster share similar textures (**Extended Data Fig. 6b**). All interest points in the remaining images in the data set were then each assigned to one cluster. The total intensity within each interest point was then counted and the fraction of intensities recorded in each interest point cluster was calculated for each cell so that each image was represented by a 100-dimensional feature vector with a sum of one.

### Non-negative tensor factorization

Each protein was represented at each mitotic stage by the average of all its vectors present at that stage. Due to binning of consecutive imaging time points, a cell can be represented by several vectors at a given mitotic stage. These duplicates were replaced by their average resulting in each cell being represented by only one vector per mitotic stage. The resulting dataset is a three-dimensional tensor **<u>X</u>** of 28 proteins *×* 100 features *×* 20 mitotic stages. We view canonical subcellular localizations as latent features of the data, that is, we assume that, at any time point, the observed vector for a protein was generated by a combination of the different canonical subcellular localizations the protein occupied at this stage. A protein vector *x* can then be expressed as the product of a subcellular localization membership vector *z* and a matrix **A** of canonical subcellular localization features: *x* = *z***A**. Therefore, we wish to model our data tensor **<u>X</u>** such that for each frontal (temporal) slice **X***_t_*,

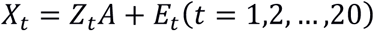

where **Z***_t_* is a matrix whose rows are localization membership vectors and **E***_t_* is a matrix containing the errors.

Given that all feature values are non-negative, a possible solution for each time point can be found by non-negative matrix factorization (NMF) of individual matrices 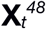. However, processing time points independently results in loss of information with the undesirable effect that different canonical localizations are learned for different time points. Simultaneous non-negative factorization of a set of matrices is a special form of non-negative tensor factorization (NTF) which can be reduced to a standard NMF using column-wise unfolding of the data tensor **<u>X</u>**^49^:

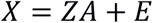

where **X** is formed by vertically stacking the **X***_t_* matrices and **Z** is formed by the correspondingly stacked **Z***_t_* matrices and **E** contains the errors. **Z** and **A** are then found using multiplicative updates^48^ to minimize the objective function ||**X** - **ZA**|| where ||.|| indicates the Frobenius norm. As a final step, the rows of **Z** are normalized to sum one. Values in **Z** can be interpreted as fractions of the amount of protein (captured by the features) present at each canonical localization.

The method requires choosing the number *k* of canonical subcellular localizations we want to represent our data with. There is no good strategy for finding this number a priori because increasing *k* corresponds to a higher resolution of the localization description e.g. a low *k* results in lumping all chromatin proteins together while a higher *k* resolves kinetochore proteins from other chromatin proteins. Thus the optimal number of subcellular localizations is partly subjective, depending on the level of granularity desired. However, we can use heuristics to help guide the choice of *k*. If the number of selected canonical localizations is too low, many proteins will share the same temporal profile, i.e. their corresponding vectors in **Z***_t_* will be highly similar for all time points. As more canonical localizations are added, we can expect more proteins to resolve into distinct profiles, i.e. the similarity between their corresponding vectors will decrease until eventually adding more canonical localizations will not improve resolution and similarity will stop decreasing. Similarity between vectors across time points can be measured using Tucker's congruence coefficient (TCC)^50^. Therefore, for each value of *k* from 2 to 25, we plot the fraction of TCC values above 0.6. The value of *k* for which the fraction of highly similar proteins reaches a low value plateau indicates that there are enough canonical localizations to describe each protein individually and therefore this value of *k* represents an upper bound on the number of canonical localizations. Following this procedure, *k* was set to seven for the current data. Because the NMF algorithm can converge to a local minimum of the objective function, ten runs with random initialization of the matrices were performed and the run with lowest value of the objective function was kept. A flattened representation of the resulting tensor can be obtained by assigning a different color to each cluster and plotting each protein distribution as a bar chart in which the height of each color band at each time point is proportional to the fraction of the protein amount in the corresponding cluster (**Extended Data Fig. 7**). A dynamic multigraph can be derived from the cluster memberships as follows: First an edge type is defined for each cluster. If two genes share a cluster at a given time point, then an edge of that type is added between them at that time point. The edge weight is set to the product of the linked genes fractions in the corresponding cluster and can be loosely interpreted as a probability of interaction. For visualization, only edges with a weight greater than an arbitrary threshold (here set to 0.3) were kept (**Fig. 4b**).

### Analysis of protein localization kinetics using supervised annotation

A multivariate linear regression model with a multivariate Gaussian response was trained with an elastic net regularization and non-negativity constraints on the coefficients with the feature vectors described above as predictors and localization vectors as response. The response vectors were defined using cells with tagged proteins known to be specific markers of unique subcellular compartments (**Extended Data Table 1**) as follows: For each of the marker proteins, the fraction of total intensity in the foreground was determined by Otsu thresholding of the 3D image stack and the corresponding protein amount assigned to the compartment with the complement assigned to cytoplasm. Each cell is thus represented by a 7-dimensional response vector containing the fraction of the tagged protein in the following compartments: chromatin, kinetochore, centrosome, spindle, midbody, nuclear envelope and cytoplasm. To deal with the compositional nature of this data, all features and response vectors are transformed using the additive log-ratio transformation^51^ with the inverse hyperbolic sine function as a generalized logarithm to handle occurrences of 0. The model with the best fit using 5-fold cross-validation was selected.

The predictions from the model were transformed back to proportions using the inverse of the log-ratio transformation then multiplied by the total number of proteins to predict the absolute number of molecules in each mitotic subcellular structure for each image. Predictions were then smoothed by local polynomial regression fitting.

To compute the anaphase dissociation kinetics for each kinetochore protein (**Extended Data Fig. 8e**), we fitted each curve between 30 min and 42 min mitotic standard time (late metaphase to telophase) with a 4 parameter sigmoidal decay function:

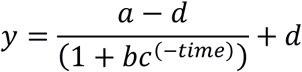

whose first and second derivatives were analytically calculated. The time of disassembly was defined as the point at which the second derivative is equal to 0 (inflection point of the curve). The disassembly rate was computed as the minimum value of the first derivative in the time interval.

### Statistics and reproducibility

For each protein, the number of cells and number of experiments that were run to collect them is reported in **Supplementary Table 1**. Unless stated otherwise, all cells for a given protein were used in the reported analyses.

### Data and code availability

All images processed in this study including original images, concentration maps, segmentation mask for both cellular and chromosomal volume and concentration maps are available in the Image Data Resource (http://idr.openmicroscopy.org^52^) under DOI: 10.17867/10000112. Further data and code are available as follows:

- All images are also available for download on the mitotic cell atlas web site http://www.mitocheck.org/mitotic_cell_atlas/downloads/v1.0.1/_cell_atlasv1.0.1_fulldata.zip (~0.5 TB).
- All source code is accessible on EMBL's GitLab instance: https://git.embl.de/grp-ellenberg/_cell_atlas and can be downloaded or cloned using the command *git clone https://git.embl.de/grp-ellenberg/mitotic_cell_atlas.git* or on the project web site at http://www.mitocheck.org/mitotic_cell_atlas/downloads/v1.0.1/mitotic_cell_atlas_v1.0.1_src.zip. Instructions to run the code are provided as a README file together with the source code. An example data set to run and test the source code can be downloaded from http://www.mitocheck.org/mitotic_cell_atlas/downloads/v1.0.1/mitoticcellatlasv1.0.1exampledata.zip.
- The data supporting the spatiotemporal mitotic cell model and the analysis is available from the mitotic cell atlas website (http://www.mitocheck.org/mitotic_cell_atlas/downloads/v1.0.1) and contains:

- Segmentation masks for the landmarks (i.e. cell boundary and chromosome mass(es)) as TIFF files (directory *mitotic_cell_model/binary_masks*).
- Snapshots of the 3D rendering of each of the spatial models in VRML and TIFF formats (directory *mitotic_cell_model/snapshots*).
- Two movies (orthogonal and oblique views) created from 3D reconstructed average landmarks (cell boundary and chromosome mass(es), directory *mitotic_cell_model/movies*).
- Average concentrations of each protein at individual mitotic stages as mat files, TIFF stacks, and tab-delimited text files (directory *protein_distributions*).
- Feature data used for the analysis (to produce **Fig. 4, Extended Data Figs. 7, 8d,e and 9**) in a tab-delimited text file (file *cell_features.txt*). This file can be used directly as input to the notebooks available in the code repository. This file also contains the mitotic standard time and stage assigned to each cell image.
- Canonical localization data (file *canonical_mitotic_clusters.h5*).
- Dynamic graph (file *dynamic_graph_adjacency_matrices.h5*).

## EXTENDED DATA LEGENDS

**Extended Data Fig. 1.**
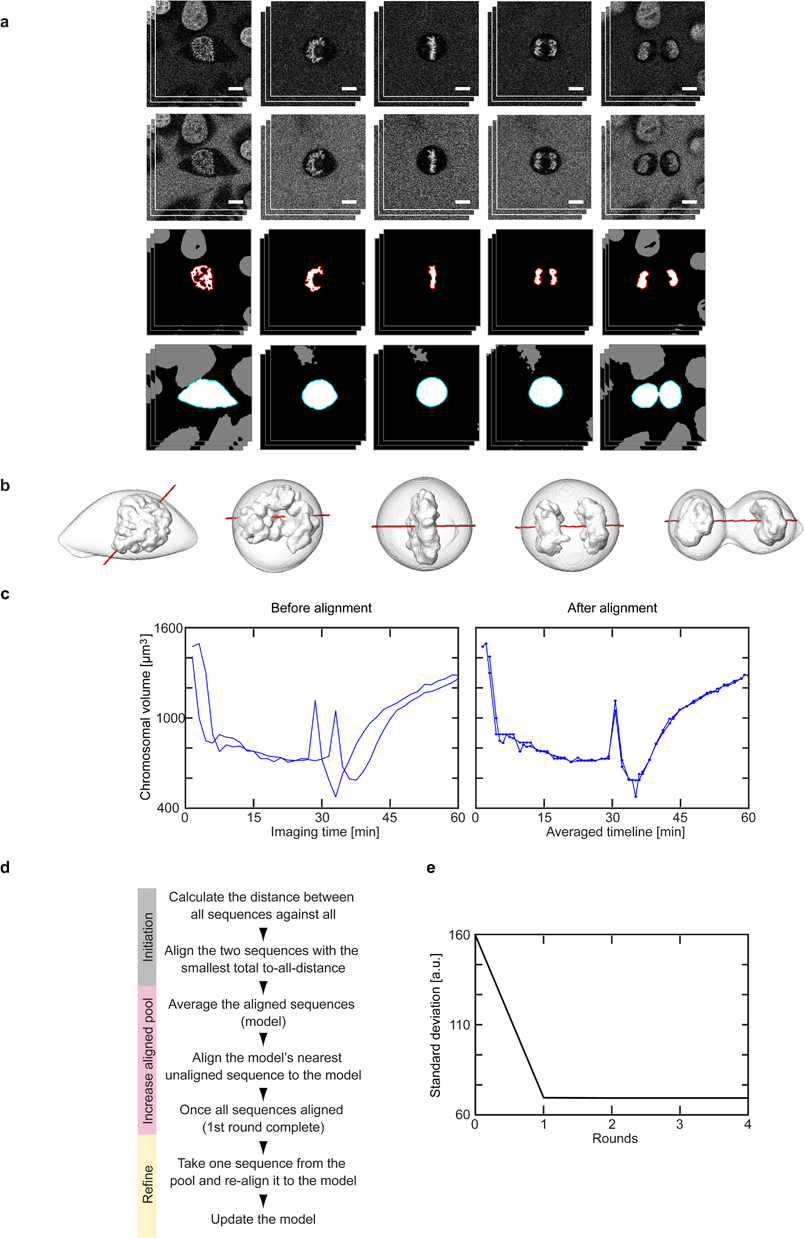
Segmentation and time alignment. (a-b) Segmentation and 3D reconstruction of landmarks. (a) Single *x-y* plane image in mCherry (587 – 621 nm, first row) and DY481XL (622 – 695 nm, second row) detection channels. Third row: detected chromatin markers where boundaries of the chromosomal volume of interest are marked in red. Fourth row: output of watershed transform on ratio image where boundary of the detected cell of interest is marked in green. Scale bar: 10 μm. (b) Reconstruction of cell and chromosomal surfaces in 3D (grey) and the predicted division axis (red). (c-e) Generating the mitotic standard time model. (c) Dynamic time warping is used to align a pair of time-resolved sequences. (d) Modified Barton-Sternberg algorithm to align 132 sequences. (e) The cumulative standard deviation of a single feature after each iteration of the algorithm. It remains nearly constant after the 2^nd^ round indicating that at termination (4^th^ round) a stable time alignment was achieved. This has been repeated 10 times and similar alignment results are obtained when the number of cells is more than 50.

**Extended Data Fig. 2.**
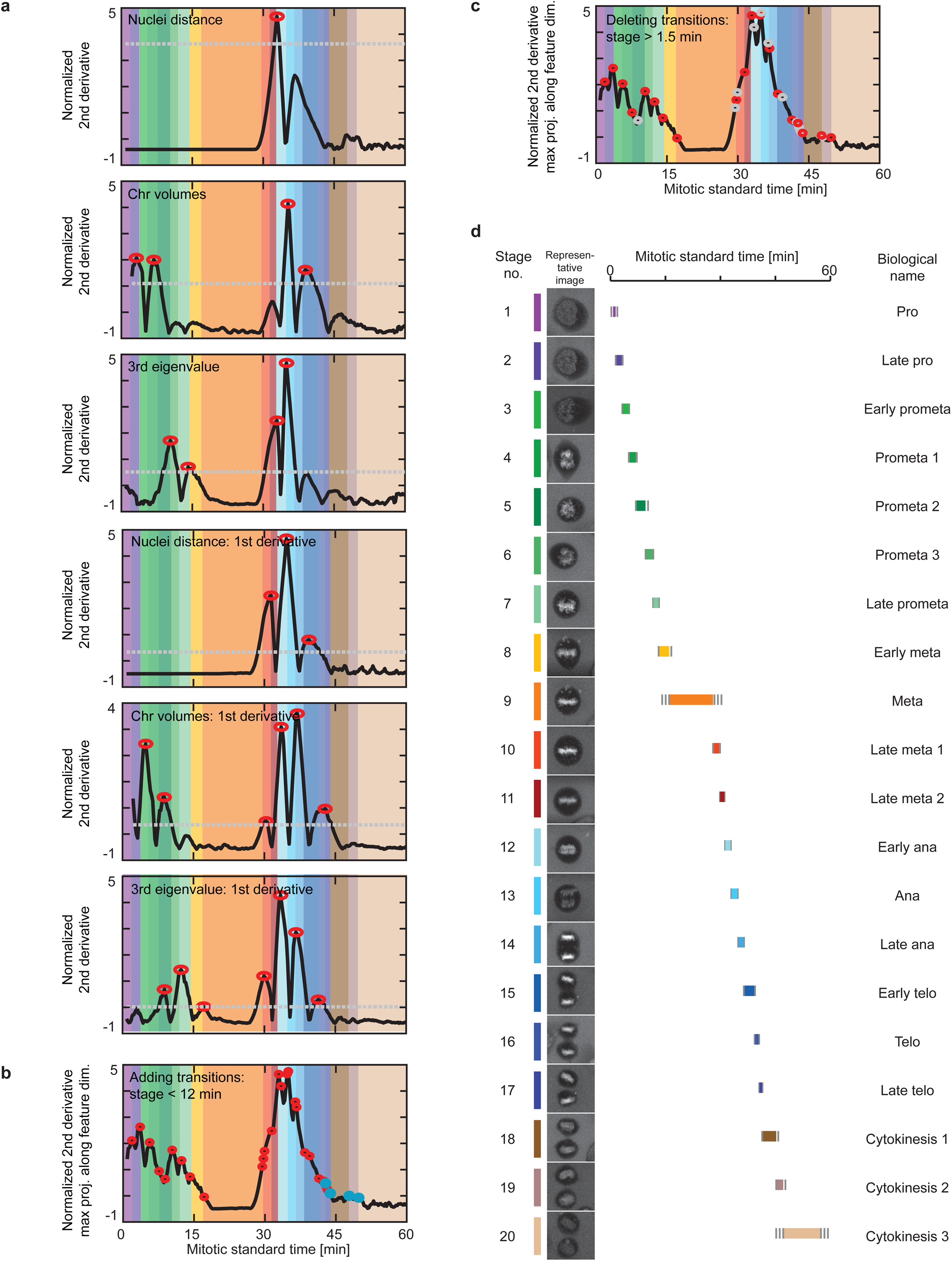
Detection of mitotic standard stages. (a) Detection of major mitotic transitions of the mitotic standard time. Peaks in the second derivatives (red circles) above a pre-defined threshold (grey lines) were detected in all feature dimensions as mitotic transitions. (b) Additional smaller peaks (blue circles) were detected to ensure a maximum duration of 12 minutes for each standard stage. (c) Transitions were deleted (grey circles) such that all stages had a minimal duration of 1.5 minutes. (d) The standard mitotic cell was represented by the cell closest to the average of each stage. Each mitotic stage was assigned duration (colored line), its duration standard deviation (grey line) and a biological annotation.

**Extended Data Fig. 3.**
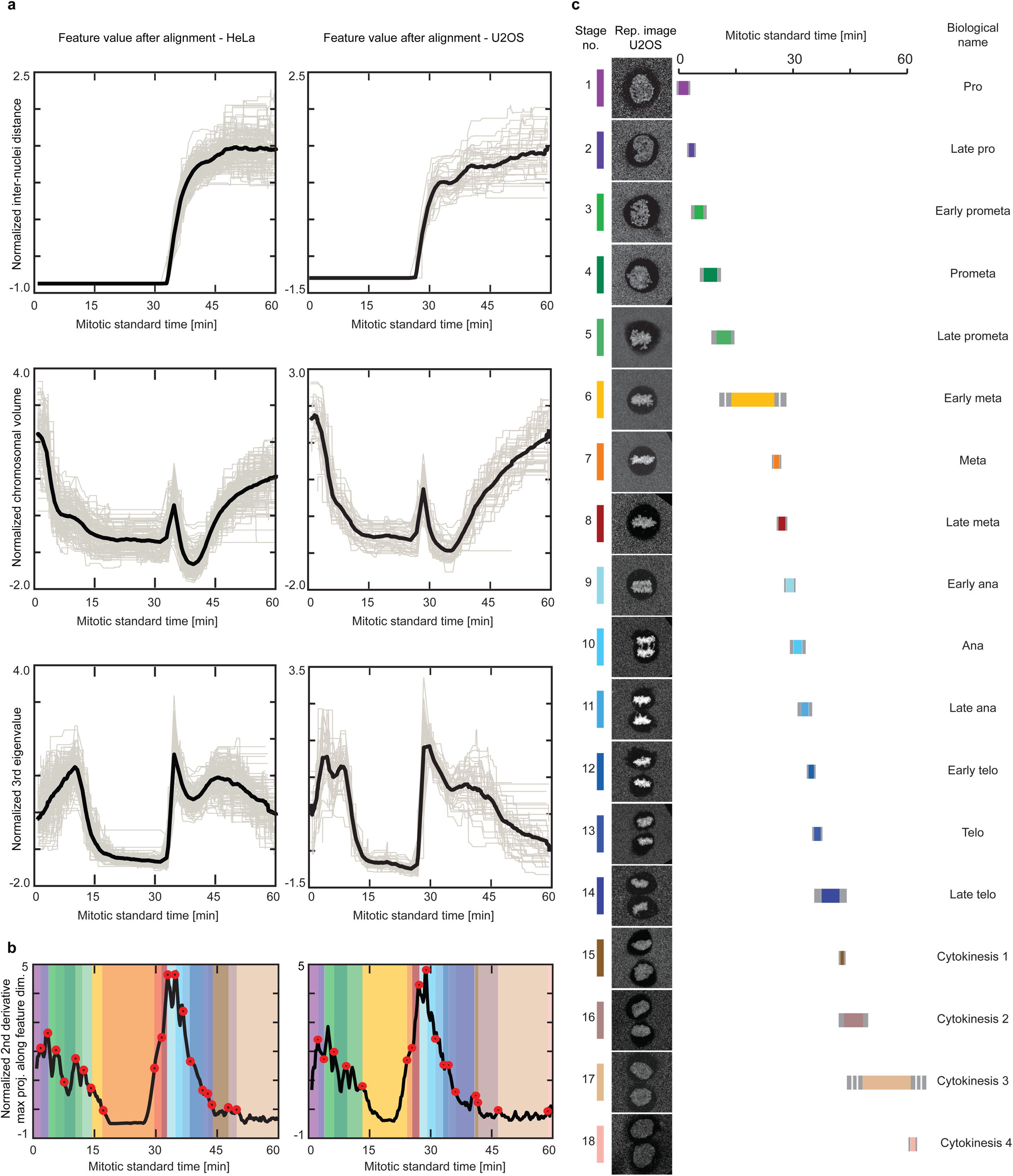
Comparison between mitotic standard time for HeLa Kyoto and U2OS cells. (a) Features used for generating the mitotic standard time model after alignment for HeLa Kyoto cells (left column) and U2OS cells (right column). Grey line: normalized feature value over time of individual cells. Black line: average. (b) Mitotic standard time transitions for HeLa cells (left panel) and U2OS cells (right panel). (c) Standard mitotic U2OS cell represented by the cell closest to the average of each mitotic standard stage. Each mitotic stage was assigned duration (colored line), its duration standard deviation (grey line) and a biological annotation.

**Extended Data Fig. 4.**
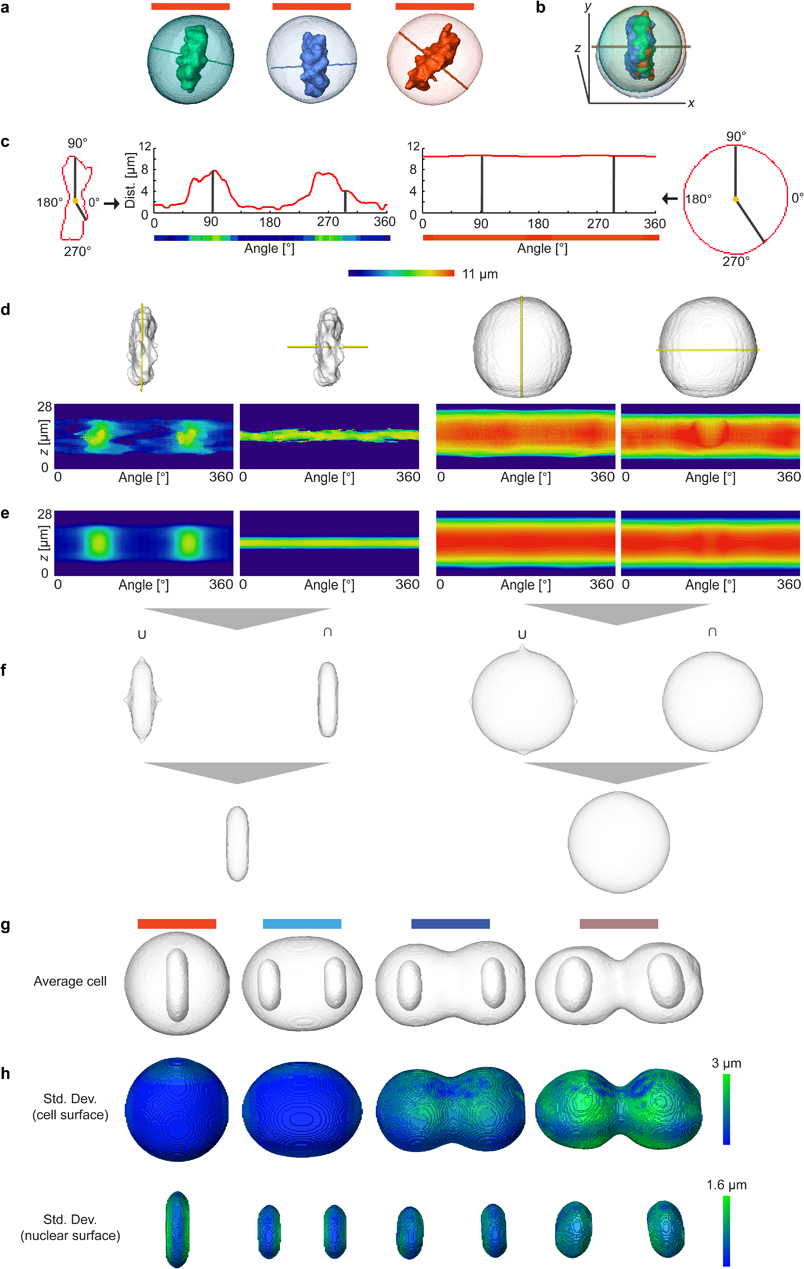
Generation of spatial model for standard mitotic stages by combining two cylindrical representations. Examples of cells in mitotic stage no. 10 (a) were registered using the predicted cell division axis as shown in (b). (c) Transformation between Cartesian and cylindrical coordinate systems. (d) Example cellular and chromosomal surfaces (grey) were transformed into the cylindrical coordinate system using two cylindrical axes (*z*-axis or predicted division axis) marked in yellow. (e) Average cellular and chromosomal surfaces in cylindrical coordinate systems. (f) Union (U) and intersection (∩) of the averaged landmarks volumes represented in the Cartesian coordinate system that were then combined to generate final cellular and chromosomal surfaces shown in the first image in (g). By averaging a large number of cells, models were generated for all mitotic standard stages with symmetrical geometries and example stages 10, 14, 16 and 19 are shown in (g). (h) The spatial variation of the mitotic standard spaces shown in (g).

**Extended Data Fig. 5.**
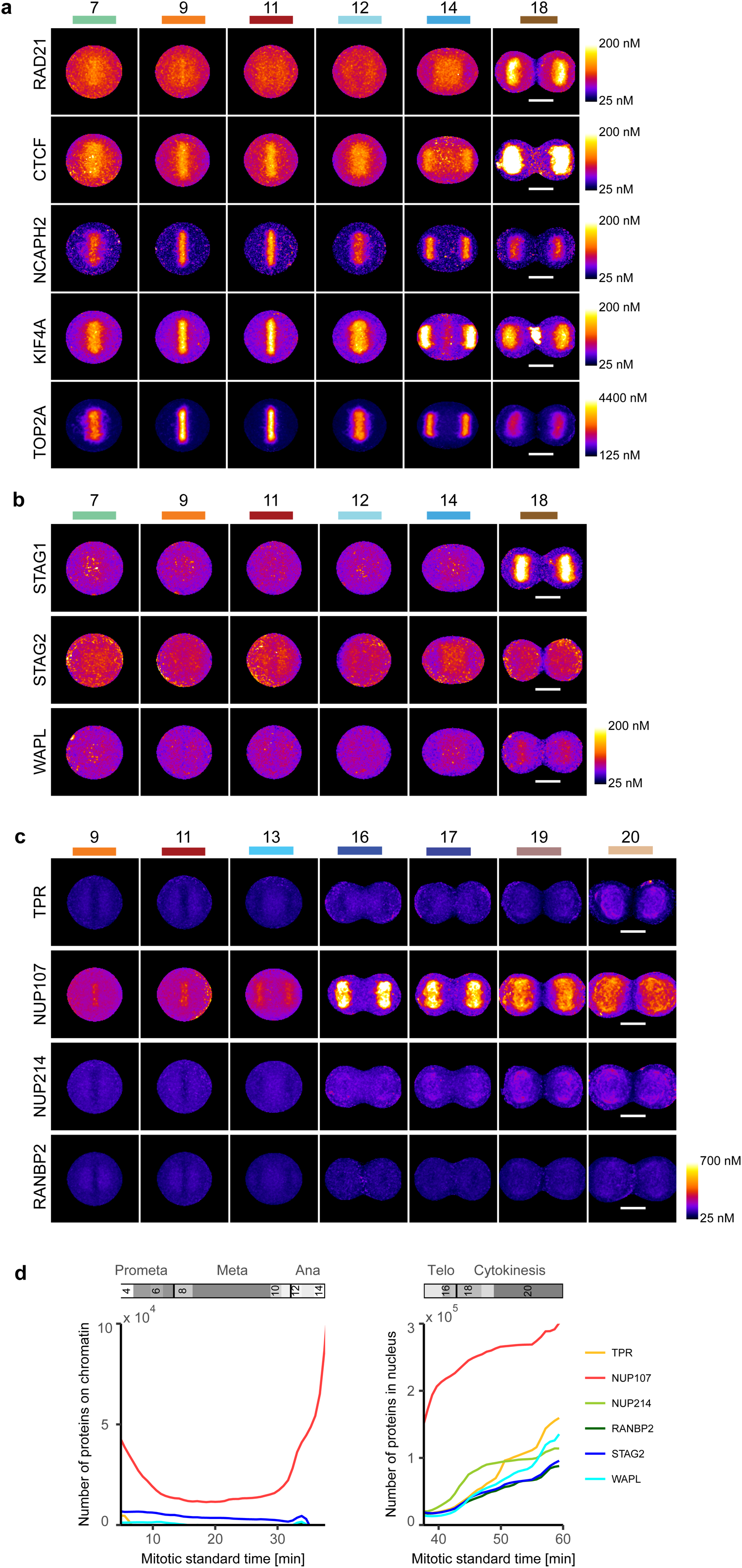
Chromatin remodelers and NUPs localization. (a-c) Maximal intensity projection from the mitotic standard model at selected stages. Scale bars: 10 μm. (a) Chromatin remodelers RAD21, CTCF, NCAPH2, KIF4A and TOP2A present on chromatin during mitosis. (b) Chromatin remodelers with weak binding to chromatin during mitosis STAG1, STAG2, and WAPL. (c) Four NUPs at selected standard mitotic stages. (d) NUPs localization as function of mitotic standard time. The curves for STAG2 and WAPL are shown as a reference and are identical to the data from **Fig. 3c**.

**Extended Data Fig. 6.**
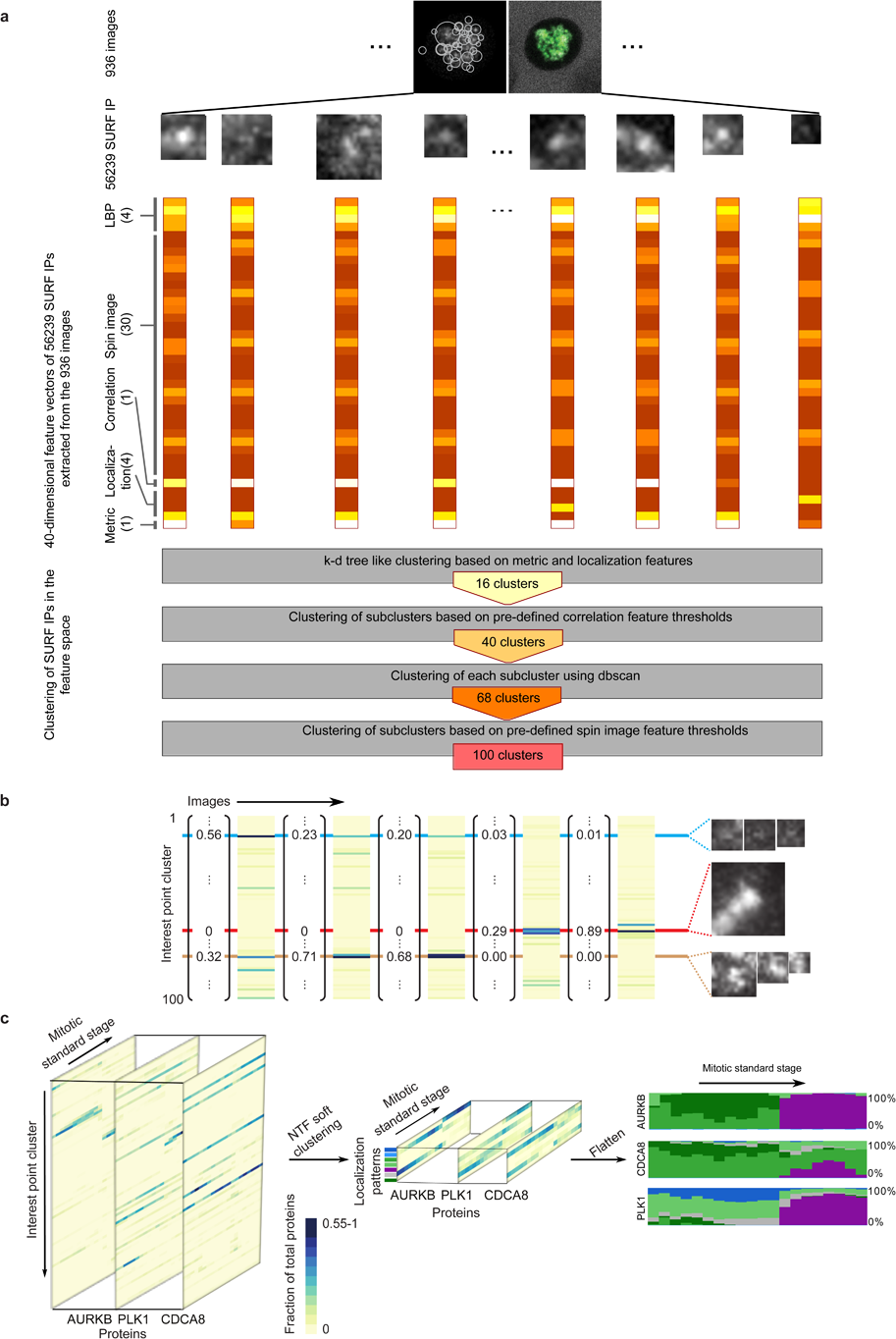
Interest point clusters and dynamic protein localization. (a) Pipeline for the definition of interest point clusters using a subset of the data. 936 images (corresponding to 5 % of the entire data set) were randomly selected from the dataset to construct a pool of interest points. Each interest point was numerically described with a 40 dimensional feature vector encoding the intensity distribution, localization and contrasts to the interest point neighborhood. Combining k-d-tree-like and thresholding-based clustering with density based clustering, the interest points were grouped into 100 clusters. (b) The remaining interest points of the data set were then assigned to the identified clusters. Thus each image was represented as the distribution of intensity in each of the 100 interest point clusters. (c) Non-negative factorization of the data tensor of proteins *×* features *×* mitotic stages (left panel) produced a non-negative tensor of reduced dimension (middle panel) whose entries can be interpreted as the fraction of protein belonging to each cluster over time (right panel, each cluster is represented by a different color and the height of a colored bar at a given mitotic stage represents the fraction of the protein in the corresponding cluster at this stage).

**Extended Data Fig. 7.**
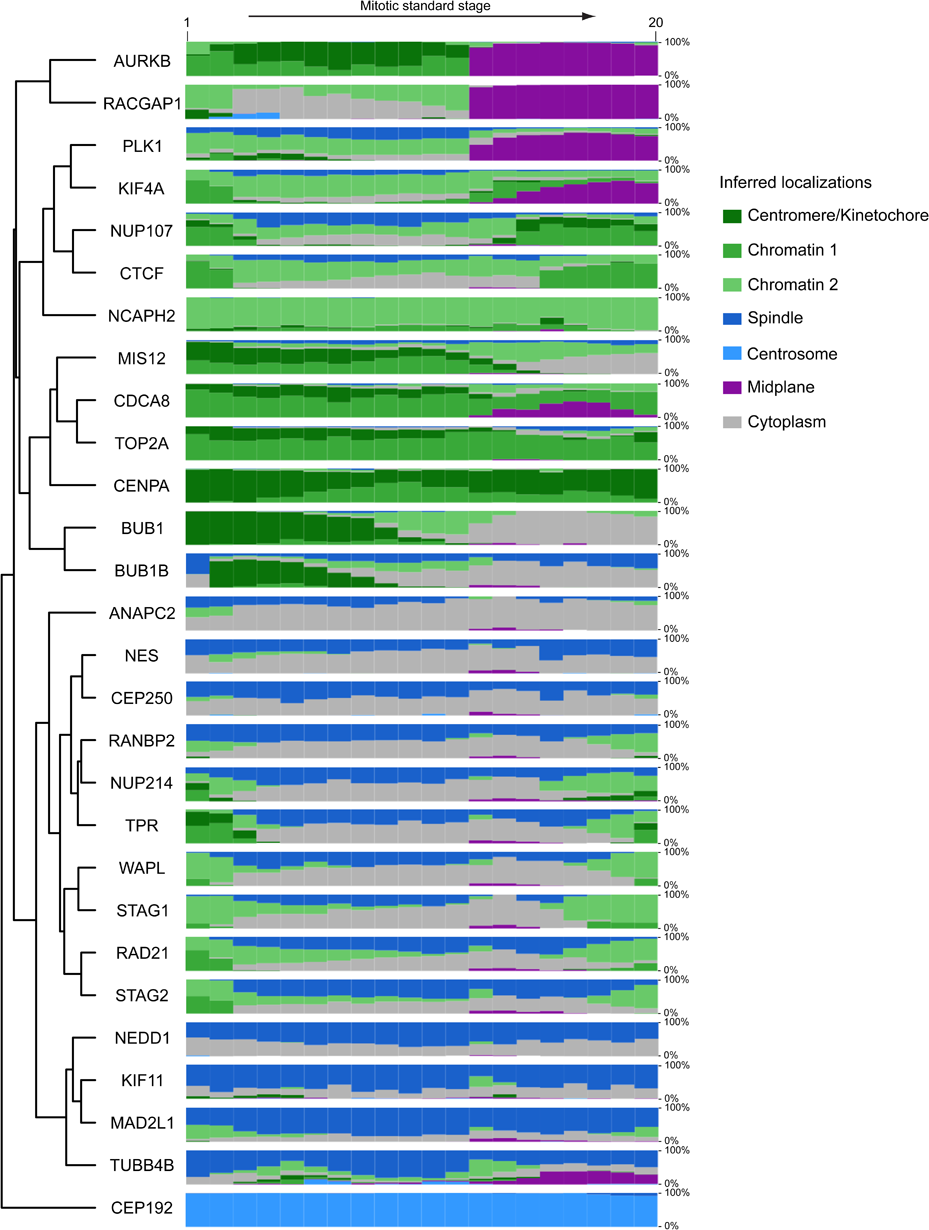
Quantitative evolution of protein subcellular localizations inferred from non-negative tensor factorization of the proteins *×* features *×* time tensor. Each subcellular localization cluster was assigned a different color and named using known information on proteins belonging to that cluster. The height of each color band at each time point is proportional to the fraction of the protein amount in the corresponding cluster at that time point. Genes were grouped by complete linkage clustering followed by optimal leaf ordering.

**Extended Data Fig. 8.**
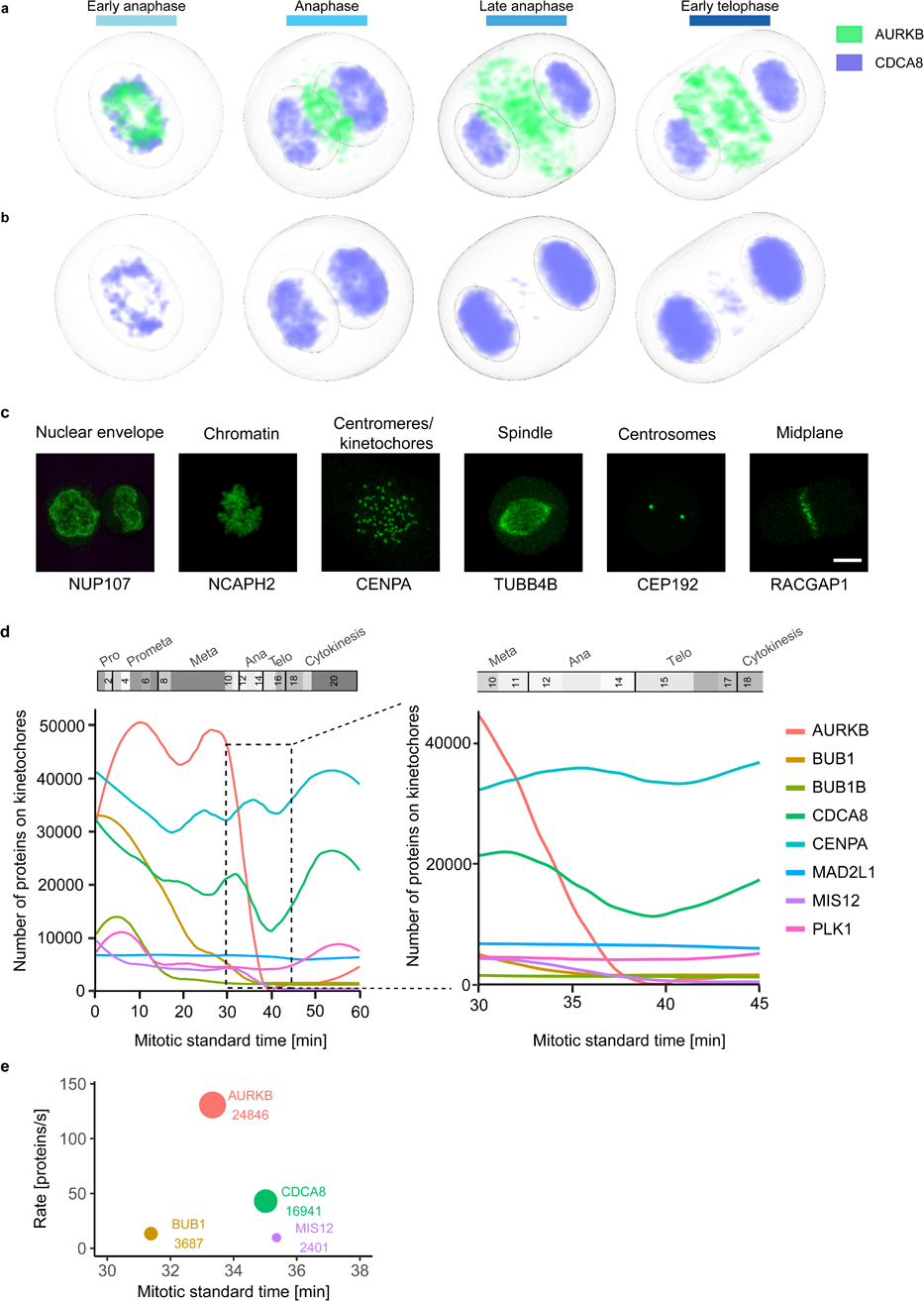
Mitotic standard model and supervised classification to investigate the dynamic localization of kinetochore proteins. (a-b) Concentration maps of chromosome passenger complex proteins AURKB and CDCA8 in anaphase and early telophase. (a) AURKB concentrates in an outer ring and a central disk. Most of CDCA8 remains on chromatin and after AURKB has already relocalized, between late anaphase and early telophase, only a small CDCA8 fraction colocalizes with AURKB in the central disk. (b) Color displaying CDCA8 was adapted to make its localization in the central disk visible. (c-e) Analyzing sub-cellular (dis)assembly kinetics using a supervised approach. (c) Example of maximally Z-projected images of marker proteins for the selected subcellular compartments used for the supervised approach. Scale bar: 10 μm. (d) Kinetics of kinetochore disassembly. The predicted number of molecules localized on kinetochore/centromeres are plotted for eight proteins in the mitotic standard time (left panel) and zoomed in for anaphase (right panel). (e) Order and rate of protein removal from the kinetochore during anaphase. The annotation and circle diameter indicate the number of molecules at the estimated average time of dissociation.

**Extended Data Fig. 9.**
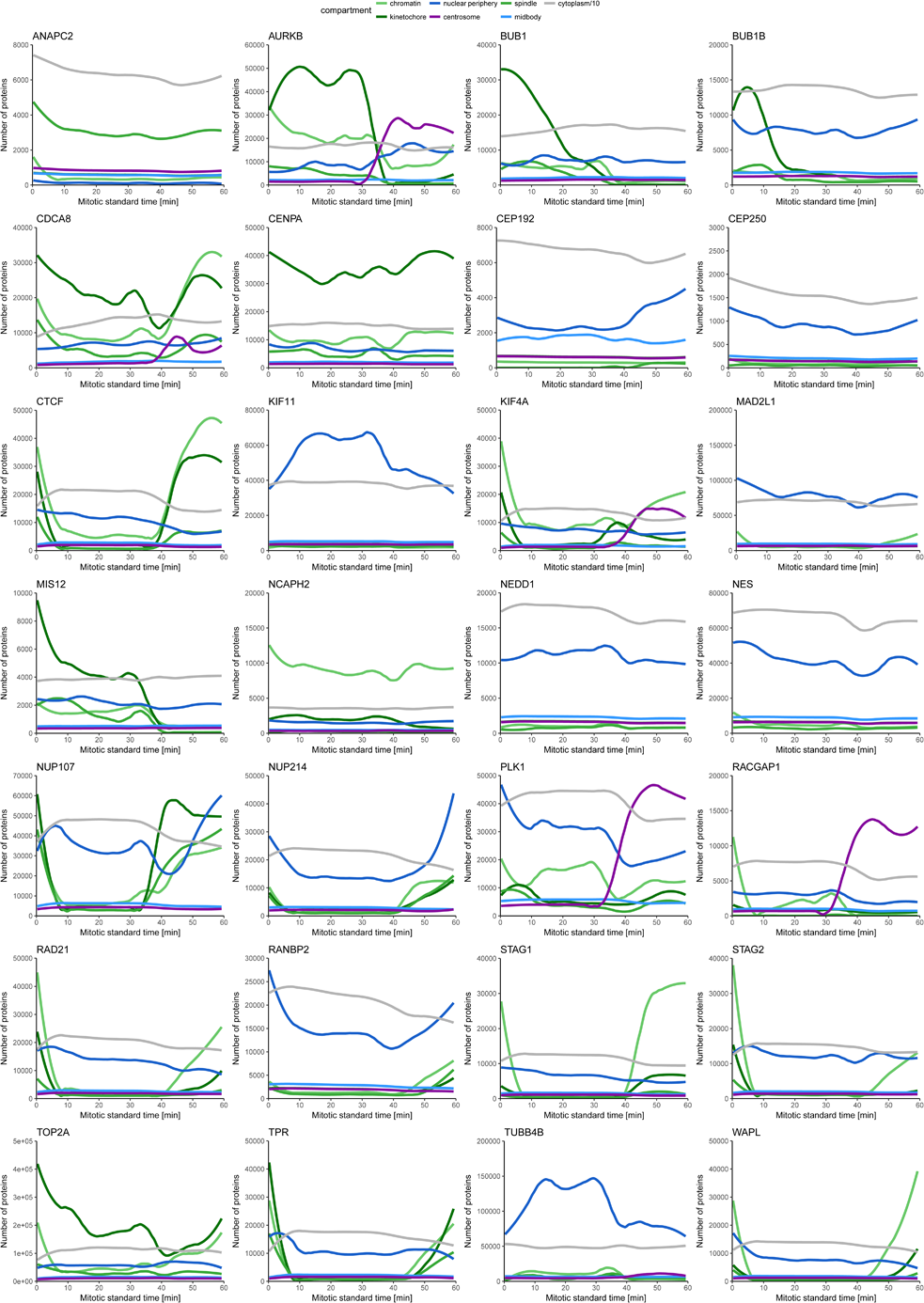
Prediction of protein molecule numbers on major mitotic subcellular structures using the supervised approach. The color scheme is adjusted to the most similar cluster identified using NTF (**Extended Data Fig. 7**). Cytoplasm values are divided by 10.

**Extended Data Table 1.** Reference structures for supervised model

| Localization | Gene | Mitotic standard stages |
| --- | --- | --- |
| Nuclear envelope | NUP107 | 15-20 |
| Chromatin | NCAPH2 | 1-20 |
| Kinetochores | CENPA | 1-20 |
| Centrosomes | CEP192 | 1-20 |
| Spindle | TUBB4B | 4-20 |
| Midbody | RACGAP1 | 12-20 |

**Supplementary Table 1.** List of cell lines used in this work

**Supplementary Table 2.** Recognition sequences and guide RNAs for genome editing

